# Shared activity patterns arising at genetic susceptibility loci reveal underlying genomic and cellular architecture of human disease

**DOI:** 10.1101/095349

**Authors:** J. Kenneth Baillie, Andrew Bretherick, Christopher S. Haley, Sara Clohisey, Alan Gray, Jeffrey Barret, Eli A. Stahl, Albert Tenesa, Robin Andersson, J. Ben Brown, Geoffrey J. Faulkner, Marina Lizio, Ulf Schaefer, Carsten Daub, Masayoshi Itoh, Naoto Kondo, Timo Lassmann, Jun Kawai, IIBDGC Consortium, FANTOM5 Consortium, Vladimir B. Bajic, Peter Heutink, Michael Rehli, Hideya Kawaji, Albin Sandelin, Harukazu Suzuki, Jack Satsangi, Christine A. Wells, Nir Hacohen, Thomas C Freeman, Yoshihide Hayashizaki, Piero Carninci, Alistair R.R. Forrest, David A. Hume

## Abstract

Genetic variants underlying complex traits, including disease susceptibility, are enriched within the transcriptional regulatory elements, promoters and enhancers. There is emerging evidence that regulatory elements associated with particular traits or diseases share patterns of transcriptional regulation. Accordingly, shared transcriptional regulation (coexpression) may help prioritise loci associated with a given trait, and help to identify the biological processes underlying it. Using cap analysis of gene expression (CAGE) profiles of promoter and enhancer-derived RNAs across 1824 human samples, we have quantified coexpression of RNAs originating from trait-associated regulatory regions using a novel analytical method (network density analysis; NDA). For most traits studied, sequence variants in regulatory regions were linked to tightly coexpressed networks that are likely to share important functional characteristics. These networks implicate particular cell types and tissues in disease pathogenesis; for example, variants associated with ulcerative colitis are linked to expression in gut tissue, whereas Crohn’s disease variants are restricted to immune cells. We show that this coexpression signal provides additional independent information for fine mapping likely causative variants. This approach identifies additional genetic variants associated with specific traits, including an association between the regulation of the OCT1 cation transporter and genetic variants underlying circulating cholesterol levels. This approach enables a deeper biological understanding of the causal basis of complex traits.

**ONE SENTENCE SUMMARY:** We discover that variants associated with a specific disease share expression profiles across tissues and cell types, enabling fine mapping and identification of new disease-associated variants, illuminating key cell types involved in disease pathogenesis.

## Introduction

Genome-wide association studies (GWAS) have considerable untapped potential to reveal new mechanisms of disease^1^. Variants associated with disease are strongly over-represented in regulatory, rather than protein-coding, sequence; this enrichment is particularly strong in promoters and enhancers^2–4^. There is emerging evidence that gene products associated with a specific disease participate in the same pathway or process^5^, and therefore share transcriptional control^6^.

We have recently shown that cell-type specific patterns of activity at multiple alternative promoters^7^ and enhancers^3^ can be identified using cap-analysis of gene expression (CAGE) to detect capped RNA transcripts, including mRNAs, lncRNAs and eRNAs^3,5^. In the FANTOM5 project, we used CAGE to locate transcription start sites at single-base resolution and quantified the activity of 267,225 regulatory regions in 1824 human samples (primary cells, tissues, and cells following various perturbations)^8^.

Unlike analysis of chromatin modifications or accessibility, the CAGE sequencing used in FANTOM5 combines extremely high resolution in three relevant dimensions: maximal spatial resolution on the genome, quantification of activity (transcript expression) over a wide dynamic range, and high biological resolution – quantifying activity in a much wider range of cell types and conditions than any previous study of regulatory variation^2,4^. Since a majority of human protein-coding genes have multiple promoters^5^ with distinct transcriptional regulation, CAGE also provides a more detailed survey of transcriptional regulation than microarray or RNAseq resources. Heritability of traits studied by GWAS is substantially enriched in these FANTOM5 promoters^9^.

Genes that are coexpressed are more likely to share common biology^10,11^. Similarly, regulatory regions that share activity patterns are more likely to contribute to the same biological pathways^5^. Transcriptional activity of regulatory elements (both promoters and enhancers^3^) is associated with variable levels of expression arising at these elements in different cell types and tissues^5^.

In order to determine whether coexpression can provide additional information to prioritise genome-wide associations that would otherwise fall below genome-wide significance, we developed network density analysis (NDA). The NDA method combines genetic signals (disease association in a GWAS) with functional signals (correlation in expression across numerous cell types and tissues, Figure 1), by mapping genetic signals onto a pairwise coexpression network of regulatory regions, and then quantifying the density of genetic signals within the network. Every regulatory region that contains a GWAS SNP is assigned a score quantifying its proximity in the network to every other regulatory region containing a GWAS SNP for that trait. We then identified specific cell types and tissues in which there is preferential activity of regulatory elements associated with selected disease-related phenotypes, thereby providing appropriate cell culture models for critical disease processes.

**Figure 1:**
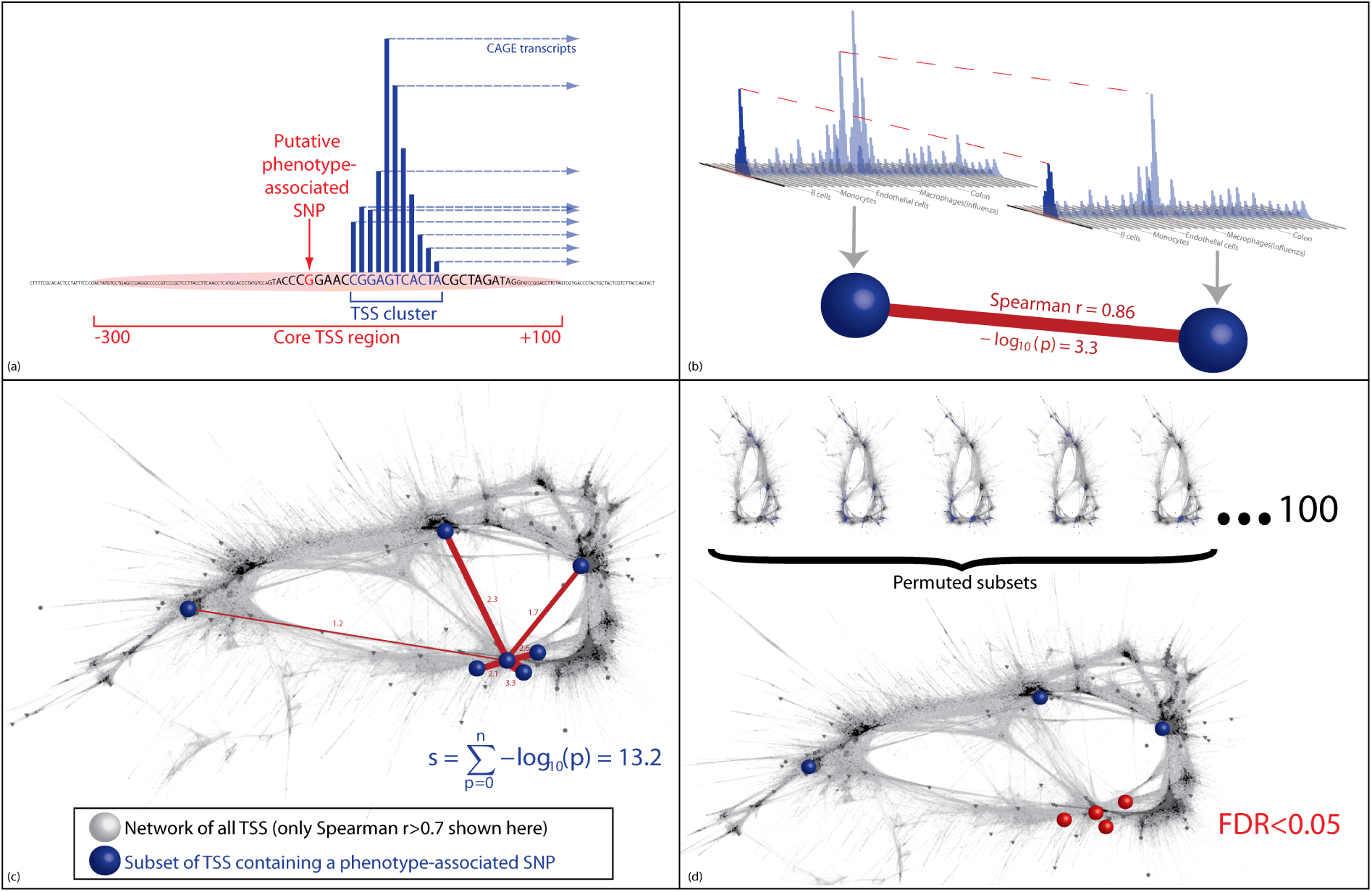
Use of NDA to detect coexpression. a) A subset of regulatory elements is identified containing disease-associated SNPs. b) The strength of the links between pairs of these regulatory regions is quantified, first as the Spearman correlation, then as the —*log*_1_0p-value quantifying the probability, specific to this regulatory region, of a Spearman correlation of at least this strength arising by chance. This is determined from the empirical distribution of correlations between this regulatory region and all other regulatory regions in the entire network of all regulatory regions in the genome. c) The subset of regulatory regions containing disease-associated SNPs form an unexpectedly dense grouping in the network, but this may not be visible in a two-dimensional representation (for illustration, this network shows all correlations between regulatory regions with Spearman *r* > 0.7, layout generated by the FMMM algorithm). The NDA score assigned to any one node is the sum of the links it shares with other nodes in the chosen subset (see Supplementary Methods for a full explanation). d) NDA scores from the input subset of regulatory elements are compared with NDA scores from permuted subsets of regulatory elements in order to quantify the false discovery rate (FDR).

## Results

### Discovery and prioritisation of GWAS hits in regulatory sequence

We defined regulatory regions as the transcription start site (TSS) -300bp and +100bp for promoters^5^, and the region between bidirectional TSS for enhancers^3^ (See Online Methods). For each of 7 GWAS studies for which high-resolution complete datasets were publicly available, we identified a set of regulatory regions containing variants with GWAS p-values below a permissive threshold (5e-8; Table 1). We devised NDA to examine the similarity in activity patterns among the set of regulatory regions detected in each GWAS (that is, the similarity in expression profile of transcripts arising from these regulatory regions).

**Table 1:** Results of coexpression analysis for a range of human traits for which high-quality data are available: Crohn’s disease, ulcerative colitis, high-density lipoprotein (HDL), low-density lipoprotein (LDL), total cholesterol, triglycerides, height, systolic blood pressure (SBP) and diastolic blood pressure (DBP). *Initial optimisation and parameterisation of the algorithm was undertaken using a random subset of data from this study.

| Trait | SNPs included, $p < 5 \times 10^{-6}$ (SNPs per million bases) | Regulatory regions containing a SNP (SNPs per million bases) | Fold enrichment for SNPs in regulatory regions | Distinct regulatory regions | Significantly co-expressed TSS ($FDR < 0.05$)(% of distinct regions) |
| --- | --- | --- | --- | --- | --- |
| Crohn's disease* | 1924 (0.6) | 133 (3.5) | 5.7 | 70 | 23 (33%) |
| Ulcerative colitis | 2162 (0.7) | 146 (3.8) | 5.5 | 83 | 20 (24%) |
| HDL | 5410 (1.7) | 260 (7.2) | 4.2 | 101 | 17 (17%) |
| LDL | 4644 (1.5) | 205 (5.2) | 3.5 | 92 | 19 (21%) |
| Total cholesterol | 6421 (2.0) | 316 (8.3) | 4.1 | 128 | 29 (23%) |
| Triglycerides | 4863 (1.5) | 254 (7.0) | 4.6 | 97 | 23 (24%) |
| Height | 8882 (2.8) | 358 (7.6) | 2.7 | 166 | 29 (17%) |
| SBP | 417 (0.1) | 20 (0.4) | 3.0 | 13 | 0 (0%) |
| DBP | 711 (0.2) | 20 (0.4) | 1.9 | 14 | 0 (0%) |

NDA detected significant coexpression (see below) among the sets of transcripts arising from regulatory regions containing variants associated with each of the following diseases and traits: ulcerative colitis, Crohn’s disease, height, HDL cholesterol, LDL cholesterol, total cholesterol and triglyceride levels (Table 1). One lower-resolution study, of blood pressure, was also analysed: in this smaller study, no coexpression signal was detected among transcripts arising near variants associated with either systolic or diastolic blood pressure (Table 1).

Significant coexpression was only detected within loci containing variants with low p-values (Fig 2a). Similar expression profiles are often seen arising from regulatory regions that are close to each other on the same chromosome, which may also span linkage disequilibrium blocks. The effect of this on the coexpression signal was mitigated by grouping nearby (within 100,000bp) regulatory regions into a single unit, unless they have notably different expression patterns (Fig 2c; Online Methods). SNPs in nearby regulatory regions are also more likely to be in linkage disequilibrium, and these regulatory regions themselves are more likely to share cis or short range trans regulatory signals in common. We checked for significant linkage disequilibrium between regulatory regions assigned to independent groups (Supplementary files 1, 4-12). At a threshold of r^2^ > 0.8, there is no linkage disequilibrium between significantly coexpressed groups; three examples of weaker linkage relationships were detected with 0.08 ≤ r^2^ ≤ 0.6 (Supplementary file 1).

**Figure 2:**
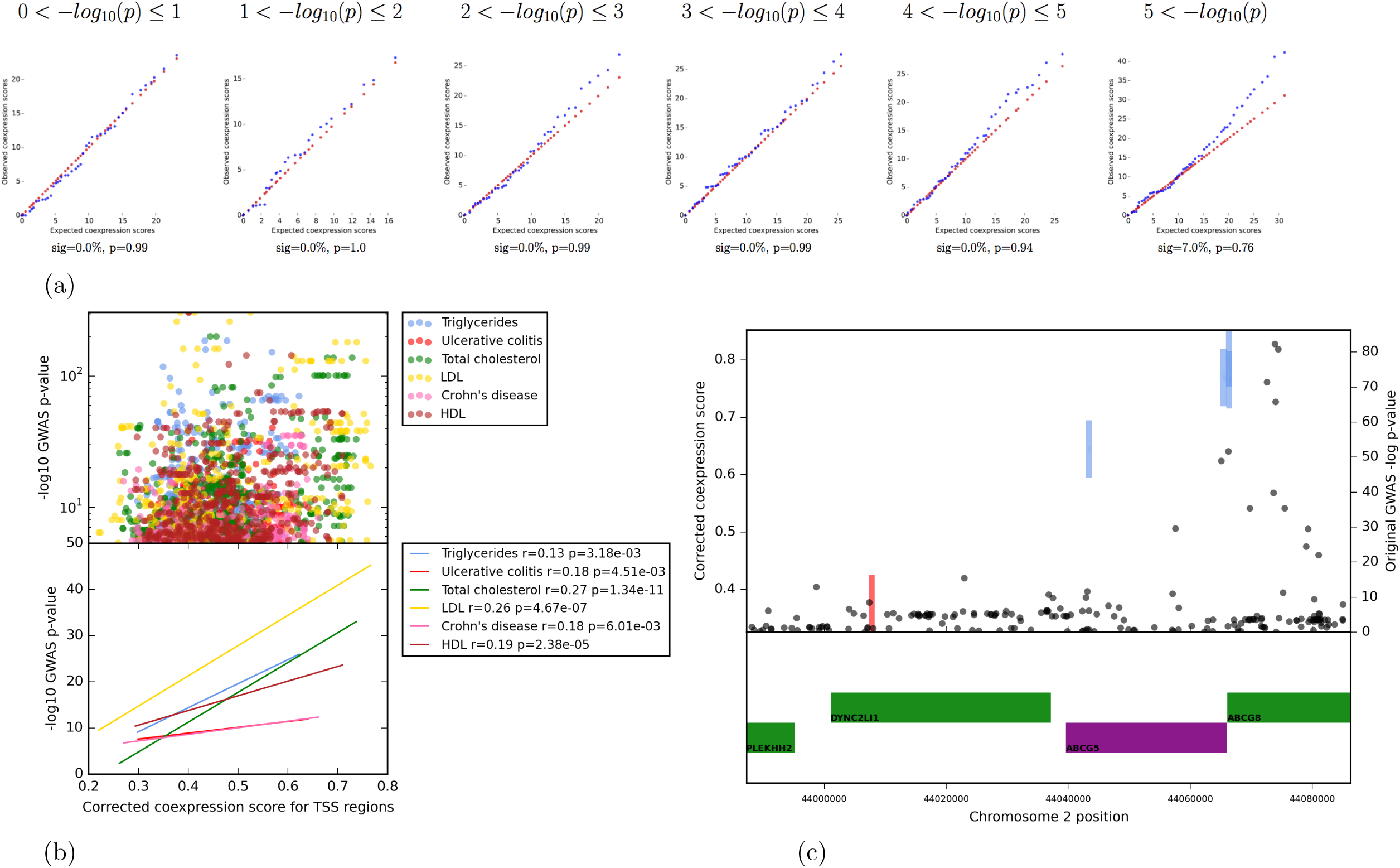
a. Change in coexpression signal in 800 SNPs selected at random from GWAS of Crohn’s disease — *log*_10_(*p*) bins from 0 to 5. No signal for coexpression is detected at weak *p*-values. Percentage of significantly coexpressed entities (hits, *FDR* < 0.05) and *p*-value (Kolmogorov-Smirnov test) comparing observed and expected distributions are shown below each plot. b. Relationship between GWAS p-value for a SNP, and coexpression scores of individual promoters assigned to that SNP. Top panel: GWAS p-values (log scale) vs corrected coexpression scores. Bottom panel: linear regression lines for data in top panel; Spearman’s r and associated p-values are shown for each trait. Only significantly coexpressed (*FDR* < 0.05) promoters are included. c. Detail of chromosomal region containing variants associated with LDL cholesterol. Top panel: Rectangles show corrected coexpression scores of individual regulatory regions; groups of regulatory regions considered as a single unit share the same colour. Black circles show GWAS *p*-values for individual SNPs. Bottom panel: known protein coding transcripts in sense (green) and antisense (purple).

Regulatory regions around individual TSS with higher coexpression scores contain variants with stronger GWAS p-values (Fig 2b), indicating that this independent signal provides additional information that may be used for fine-mapping causative loci (Fig 2c).

In order to enable the detection of new regulatory regions with strong coexpression relationships, we chose a permissive p-value threshold for trait association of 5×10^-6^(see Online Methods). GWAS data for Crohn’s disease^12^ were used for initial optimisation of the NDA approach; among GWAS datasets for phenotypes that were not used in algorithm development (i.e. all apart from Crohn’s disease), 0-24% of regulatory regions containing a GWAS SNP showed significant coexpression with other regulatory elements associated with the same phenotype (FDR < 0.05, compared with 100 permuted subsets of equal size; see Online Methods).

For a given disease, regulatory regions containing GWAS variants are coexpressed if they share similar activity patterns (i.e. similar expression patterns among transcripts arising from these regulatory regions) with other regulatory regions implicated in that disease. Figure 3 shows significant coexpression superimposed on a two-dimensional representation of the entire network of pairwise correlations. Since activity (transcript expression) was measured in numerous samples, the true proximity of regulatory regions to one another cannot be accurately represented in two dimensions – a perfect representation would require as many dimensions as there are unique samples. However, the NDA method is designed to quantify proximity in network space, so that significantly coexpressed elements are detected, even if they are not directly adjacent on a twodimensional representation of the network (Figure 3). Among strong coexpression was seen between loci that were widely separated on the genome (Figure 4).

**Figure 3:**
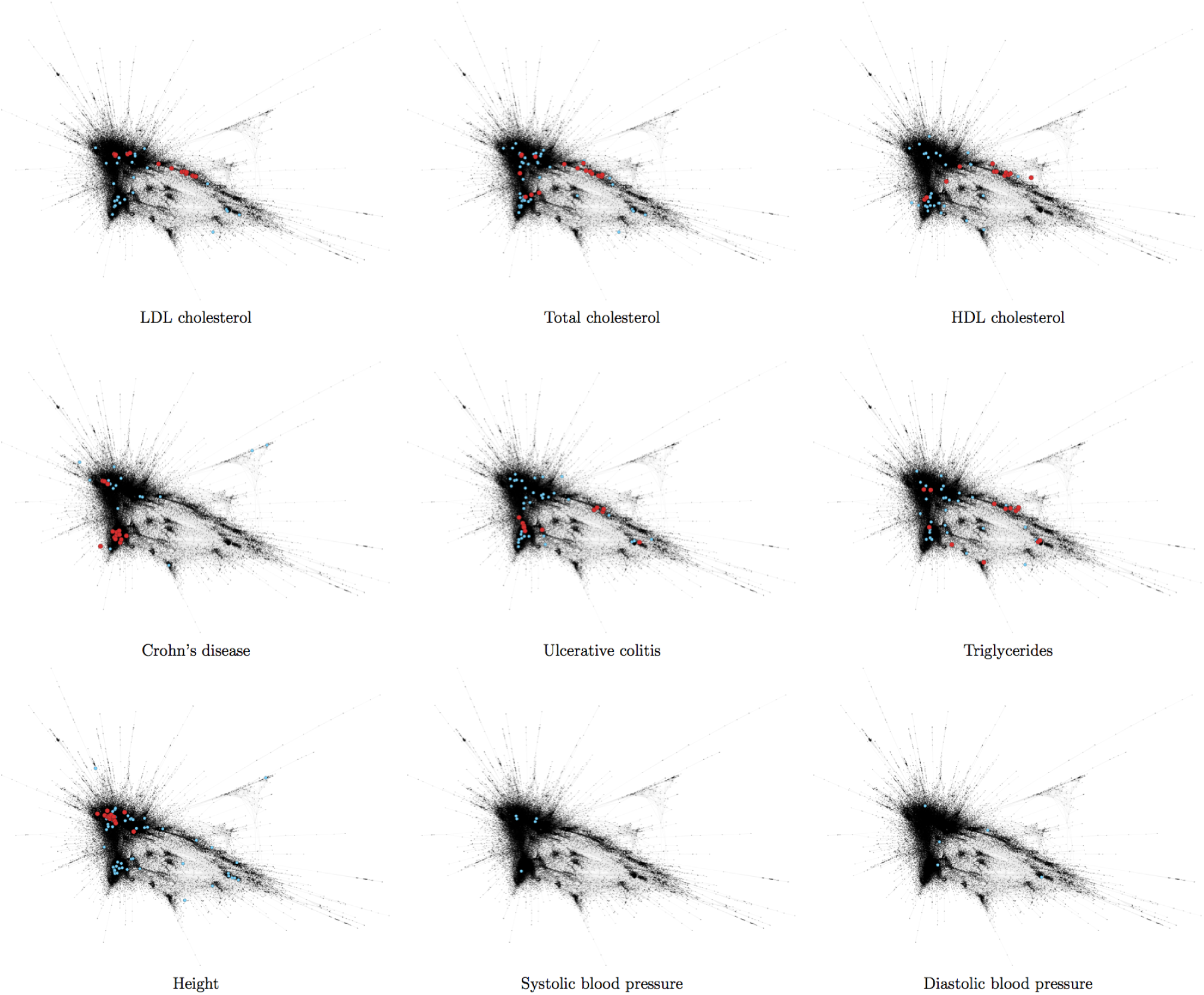
Network layouts (Spearman r > 0.5, FMMM layout algorithm, largest component only is shown) showing position of significant hits on a two-dimensional network representation of FANTOM5 regulatory regions. Red circles: significantly-coexpressed (*FDR* < 0.05) regulatory regions containing a putative GWAS hit (*p* < 5 × 10^-6^) for this trait. Blue circles: regulatory regions containing a putative GWAS hit (*p* < 5 × 10^-6^) for this trait that are not significantly coexpressed (*FDR* > 0.05).

**Figure 4:**
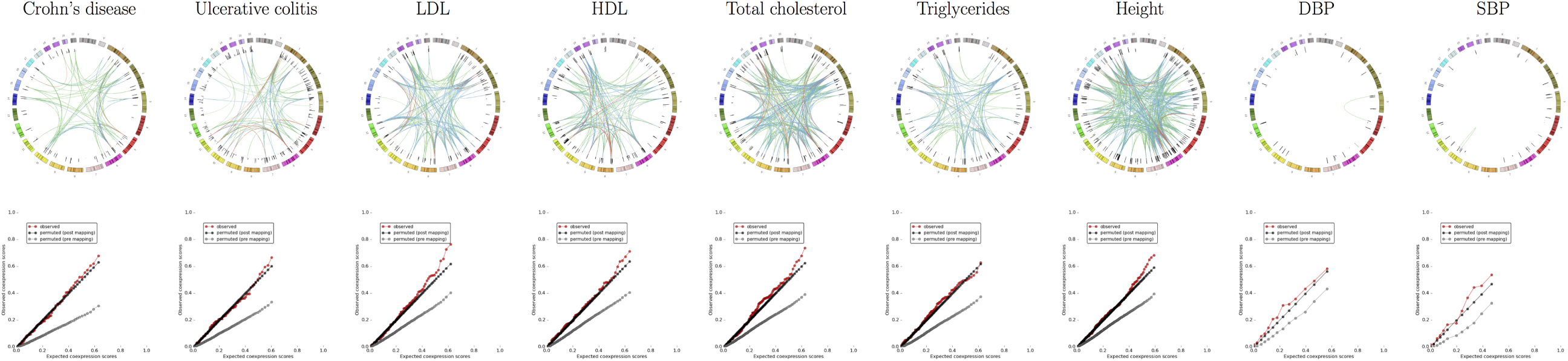
(Top panels) Circular plots of coexpression links between different locations on the genome, illustrating the spatial separation of highly-correlated regulatory regions. The coloured outer circle shows an end-to-end concatenated view of the human chromosomes. The black inner circle shows *log_10_* GWAS p-values for included SNPs. Links depict an association between two regulatory regions containing these wSNPs and are coloured according to — *log*_10_(*p*)(line colour indicates *log*_10_(*p*): red>3, blue> 2, green> 1.5). (Bottom panels) Quantile-quantile plots showing observed and expected coexpression scores. Expected coexpression scores are derived from circular permuted subsets of regulatory regions (post-mapping permutations; black circles) or SNPs chosen by circular permutations against the background of all SNPs genotyped in each study. Data are shown for Crohn’s disease, ulcerative colitis, high-density lipoprotein (HDL), low-density lipoprotein (LDL), total cholesterol, triglycerides, height, systolic blood pressure (SBP) and diastolic blood pressure (DBP)

The coexpression signal essentially combines the signal for association in a GWAS with the location and activity pattern of regulatory regions on the genome. We deliberately chose a permissive GWAS p-value threshold in order to enable the detection of new signals that did not achieve genome-wide significance in the original studies. For example, we found that coexpressed transcripts for both LDL and total cholesterol (TC) arise from promoters for well-studied genes such as APOB^13^ and ABCG5^14^, but also from regulatory regions not previously associated with cholesterol levels. A promoter for SLC22A1, which encodes an organic cation transporter, OCT1^15^, is strongly coexpressed among elements associated with both conditions (Supplementary File 1). OCT1 transcription is regulated by cholesterol^16^ and the transporter regulates hepatic steatosis through its role in thiamine transport^17^. This action of OCT1 is inhibited by metformin^17^, an oral hypoglycaemic agent whose cholesterol-lowering effect^18^ is not well understood^19^. Full results of coexpression analyses are in Supplementary File 1, and online at www.coexpression.net.

### Cell-type and tissue specificity

The significantly-coexpressed networks detected here could be regarded as revealing the signature expression profile, at least within the FANTOM5 dataset, for a given disease or trait. We next explored whether these signature expression patterns reveal cell types or biological processes that may contribute to the trait or disease susceptibility.

We therefore ranked cell types and tissues by transcriptional activity for each of the significantly-coexpressed loci for each trait, and combined the rankings using a robust rank aggregation^20^ (Online Methods). By first detecting the characteristic expression signature associated with a given phenotype using only high-resolution GWAS data, and then detecting the cell type and tissue activity profiles that underlie this signature, we improve on the statistical power of previous methods that have attempted to detect cell-type specific signatures of disease^4,6,21^. Strong signals reported previously are highly significant in our analysis; for example genetic loci associated with cholesterol are transcriptionally active in hepatocytes and liver tissue^6^(Supplementary File 8).

This analysis reveals robust cell-type associations that have important implications for understanding disease pathogenesis. For example, cell-type associations with Crohn’s disease were restricted to immune cells, particularly monocytes exposed to inflammatory stimuli (Supplementary File 4). In contrast, cell type associations with ulcerative colitis were statistically significant in rectum, colon and intestine samples, and in a distinct group of immune cells: macrophages exposed to bacterial lipopolysaccharide (Supplementary File 5). This is consistent with the view that ulcerative colitis, in which disease processes are primarily restricted to the colon and rectum, is a consequence of dysregulation of processes that are intrinsic to the large bowel, including epithelial barrier function^22^, whereas Crohn’s disease is a multisystem autoimmune disorder with more diverse extra-intestinal manifestations^23^, consistent with a primary immune aetiology.

## Discussion

The development of high-throughput genotyping methods has led to an explosion of associations between genetic markers and human diseases^24^. The results presented here are a step towards overcoming the next challenge for this field: making sense of these associations to advance the practice of medicine. There has been increasing recognition of the potential to utilise prior knowledge to improve detection and interpretation of genome-wide signals^25^. The results of our analysis demonstrate that there is biological information in the coexpression of genetic variants associated with a particular disease that can provide the basis for prioritising variants that would not otherwise meet standard thresholds for genome-wide statistical significance.

We report relationships between numerous regulatory regions that are not associated with named genes – a restriction that has previously limited the transition from genetic discovery to biological understanding^26-30^. The analysis reveals the impact of specific enhancers and promoters that may be remote from the genes they regulate, or may contribute to tissue-specific regulation of a gene that may otherwise appear to be more widely-expressed.

Even for those disease-associated variants that can be reliably assigned to a named gene, previous attempts to draw functional inferences have, by necessity, relied on published data^26^,annotated biological pathways^31^, or gene sets^30,32^. Although many important insights have been gained from these approaches, they share a fundamental limitation: reliance on existing knowledge. This restricts the ability to exploit the potential of genomics to deliver insights into new, previously unseen, mechanisms of disease^33^.

The data used for development and testing of the coexpression approach were from large meta-analyses that incorporate genotyping (or imputation) of genetic variants at extremely high resolution, increasing the probability that variants will be found within regulatory regions. In future, the availability of whole-genome sequencing can reasonably be expected to produce many additional high-quality datasets for coexpression analysis. In principle, the NDA approach can be generalised to any network in which it is desirable to quantify the proximity of a subset of nodes.

The scale, depth and breadth of the FANTOM5 expression atlas, together with the NDA approach, enable detection of subtle coexpression signals for regulatory regions that have previously been undetectable. As additional genetic studies become available at greater genotyping resolution, we anticipate that this method will detect new genetic associations with disease, coexpressed modules underlying pathogenesis, identify critical cell types implicated in mechanisms of disease.

## DATA ACCESS

The FANTOM5 atlas is accessible from http://fantom.gsc.riken.jp/data/

An online service running the coexpression method is available at http://coexpression.net

## ACKNOWLEDGEMENTS

We would like to express our gratitude for the diligence and professionalism of the entire FANTOM5 consortium and to the members of the IIBDGC group, GIANT consortium, and Global Lipids consortium for freely sharing their data. We are particularly grateful to the tens of thousands of patients and healthy volunteers who donated DNA and other material to these studies.

JKB gratefully acknowledges funding support from a Wellcome Trust Intermediate Clinical Fellowship (103258/Z/13/Z) and a Wellcome-Beit Prize (103258/Z/13/A), BBSRC Institute Strategic Programme Grant to the Roslin Institute (BBS/E/D/20241864), the UK Intensive Care Foundation, and the Edinburgh Clinical Academic Track (ECAT) scheme. Funds were provided to the Roslin Institute through a BBSRC Strategic Programme Grant (JKB, SC, CSH, GJF, TCF, DAH; BBS/E/D/20211551, BBS/E/D/20231760). We acknowledge the financial support provided by the MRC-HGU Core Fund (CSH, AT). FANTOM5 was made possible by a Research Grant for RIKEN Omics Science Center from MEXT to YH and a Grant of the Innovative Cell Biology by Innovative Technology (Cell Innovation Program) from the MEXT, Japan to YH. RIKEN Centre for Life Science Technologies, Division of Genomic Technologies members (RIKEN CLST (DGT)) are supported by institutional funds from the MEXT, Japan. ARRF is supported by a Senior Cancer Research Fellowship from the Cancer Research Trust and funds raised by the Ride to Conquer Cancer. JCB is supported by Wellcome Trust grant WT098051. GJF acknowledges the support of an NHMRC Career Development Fellowship (GNT1045237), NHMRC Project Grants (GNT1042449, GNT1045991, GNT1067983 and GNT1068789), and the EU FP7 under grant agreement No. 259743 underpinning the MODHEP consortium. MR was supported by grants from the Deutsch Forschungsgemeinschaft, the German Cancer Aid and the Rudolf Bartling Foundation. RA was supported by funding from the European Research Council (ERC) under the European Union’s Horizon 2020 research and innovation programme (grant agreement No 638273). US and VBB are supported by the KAUST Base Research Fund to VBB and KAUST CBRC Base Fund. RMP is supported by grants from the US National Institutes of Health (R01-AR057108, R01-AR056768, U01-GM092691 and R01-AR059648) and holds a Career Award for Medical Scientists from the Burroughs Wellcome Fund. RA and AS were supported by funds from FP7/2007-2013/ERC grant agreement 204135, the Novo Nordisk foundation, and the Lundbeck Foundation and the Danish Cancer Society. CAW is supported by a Queensland Government Smart Futures Fellowship, and samples were collected under Australian National Health and Medical Research council project grants 455947 and 597452, under agreement from the Australian Red Cross 11-02QLD-10 and the University of QLD ethics committee.

## Authors’ contributions

JKB conceived the study, designed and led the analyses and wrote the manuscript. AB and AG contributed to computational optimisation and description of methods. AB and SC generated network and circos images, respectively. CH, JB, JBB, TF, and AT advised on statistical and network analysis methods. ML managed the data collection, including annotation, expression profiling, metadata association and archiving. CW, JS, NH and TF contributed biological expertise. CW, RA, JS, AS, MR, VB, and PH advised on methodology. ARRF, MI, CD, NK, TL, JK, HS, HK, YH, and PC organised the FANTOM5 project including sample collection, data production, mapping and tag clustering. JKB, ARRF, DAH, TF, GJF, PC and YH provided resources. DAH and ARRF advised on methodology, and contributed to the manuscript. All authors contributed to and approved the final version of the manuscript.

## SUPPLEMENTARY METHODS

### 1 Regulatory regions

For the purpose of this analysis, promoters identified in the FANTOM5 dataset were defined as the region from -300 bases to +100 bases from a transcription start site (Figure 1a, main paper). Previous analysis demonstrated that this covers the areas of maximal sequence conservation across species[4] and the core region of transcription factor binding[12]. Since eRNA TSS are considerably longer than promoter TSS (median length(IQR) 272(173-367) vs 15(9-26)), enhancers were defined by the range covered by eRNA transcription start sites[10].

### 2 Coexpression algorithm

For each GWAS study, SNPs were identified that lie within either a functional promoter or enhancer. Any promoter or enhancer that contained a variant putatively associated with a given phenotype was considered to be candidate phenotype-associated regulatory region. A pairwise coexpression matrix was then generated across the full FANTOM5 dataset of promoters and enhancers, in which each node is a regulatory region, and edges reflect the similarity in activity (expression) patterns arising at these regulatory regions, across different cell types and tissues.

To test the hypothesis that regulatory regions genetically associated with a given phenotype are more likely to be coexpressed, we devised a method to quantify coexpression among a pool of putative phenotype-associated regulatory regions (network density analysis; NDA). This approach avoids arbitrary cut-offs between clusters (or communities) of nodes, and yields a single value for each node, quantifying the closeness with all other nodes in a particular subset (network density). NDA was used to integrate the putative association between a regulatory sequence and the phenotype of interest (indicated by the presence of a phenotype-associated SNP), with the coexpression similarity between this node with other nodes that are also putatively associated with the same phenotype.

#### 2.1 Principle of network density analysis (NDA)

NDA integrates information from two distinct and independent sources: the relationships between nodes in the network, and the choice of subset. In the present work, nodes are regulatory regions, the subset is those regulatory regions that contain variants associated with a particular phenotype, and the relationships are Spearman’s rank correlations. However, the NDA approach is generalisable to any network of pairwise relationships.

Within a network of all possible pairwise relationships between nodes, a subset of nodes is selected that share a particular characteristic.

Within this subset of nodes, every pair of nodes is considered. Each relationship between two nodes is expressed as the — log_10_ of the empirical probability of a relationship at least as strong occurring between the chosen node and another, randomly-chosen, node from anywhere in the whole network. These probabilities are specific to each node and are directional. The NDA score is the sum of the −log_10_(*p*) values for a node in the chosen subset and all other nodes within the subset. The NDA score therefore quantifies the density of this subset of nodes in network space. The purpose of using the empirical probability of a correlation, rather than the raw correlation metric, is to control for bias in favour of highly-connected nodes, as would occur if one expression profile were very common. Finally, the NDA score is assigned its own *p*-value by comparison to that obtained using randomly permuted subsets (see below). If the network contains no additional information about this subset of nodes, then the relationships between nodes in the chosen subset will be no stronger than the relationships seen in permuted subsets.

#### 2.2 Application to coexpression of regulatory regions

From the set of all nodes in a network, a subset is selected because they share some characteristic. In the case of the genomic analyses reported here, the nodes are TSS, and the subset of interest is those TSS that contain a variant that has some evidence of association with a particular trait. Throughout this paper, we have defined the set of phenotype-associated transcription start sites, *R*, as follows: the set of regulatory elements associated with phenotype-associated single nucleotide polymorphism within 300bp (promoters) or 0bp (enhancers) upstream from a FANTOM5 transcription start site (TSS) and 100bp (promoters) or 0bp (enhancers) down-stream. In order to enable the detection of new associations, we use a deliberately permissive threshold. We define as “putatively-significant” a SNP-phenotype association of *p* < 5 × 10^−6^. Let the integer variable *i* be used to index the base pairs (bp) of the genome. For a given trait, the set of input SNPs, *K*, are those that have a putatively-significant association with that trait at our chosen threshold. If we let *TSS_start_* equal the base pair index 300bp (promoters) or 0bp (enhancers) upstream from a FANTOM5 transcription start site (TSS) and *TSS_end_* 100bp (promoters) or 0bp (enhancers) downstream, the set, *P*, of putative trait-associated promoters is given by:

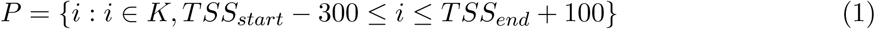

and the set *E* of enhancers containing a putative trait-associated SNP is given by:

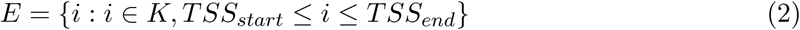

giving a total set of regulatory regions:

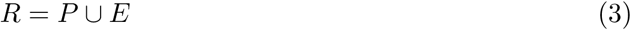

#### 2.3 Linkage disequilibrium (LD) - grouping nearby regulatory regions

Input SNPs from GWAS results tend to be in LD with nearby variants. There is therefore a risk of spurious coexpression, since nearby regulatory regions are also likely to share regulatory influences, such as chromatin accessibility, enhancers, and lncRNAs. One solution to this would be to filter input SNPs by LD. However this would require that LD relationships for all SNPs be known for all of the populations from which SNP association data were derived, which is not the case. It would also risk removing functionally important regulatory regions from the analysis, by choosing only one SNP per LD block.

In order to overcome these problems, we sought to identify those regulatory region-associated SNPs within a given region that are most likely to contribute to a given subnetwork of putative phenotype-associated regulatory regions. By the definitions described above, these will be those regulatory regions with the highest NDA score. Regulatory regions are considered for combination if they are separated by 100,000bp or less. If any regulatory regions within this range has a correlation *p*-value of less than 0.1 with any other regulatory regions in the range, they are combined. A single representative regulatory region is then chosen - the regulatory region with the largest NDA score in the group, derived from a network comprised of all other groups.

In order to confirm that spuriously significant coexpression signals are not being generated because of LD, we used the ENSEMBL Perl API for the 1000 genomes phase 3 data (CEU) to search for variants in LD with each SNP lying within the chosen regulatory region for each group. Variants in LD with a variant in any other chosen regulatory region are reported.

#### 2.4 Coexpression matrix

Let *A* be the set of all nodes in the whole network. Each member of *A* is a node in an interaction network. For each *i* ∈ *R*, Spearman’s rank correlation, *x*, is calculated with each other node in *R*. The probability, *p*, of a correlation as strong or stronger as the index correlation, *x*, arising by a chance pairing between the index node and any other node (*n*_(*r*>*x*)_) is inferred from the empirical distribution of all correlations (*r*) of the index node in *A*.

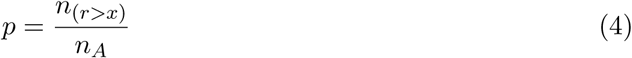

#### 2.5 Network density analysis

For every node in the set *R*, a score *s* is calculated to summarise the strength of interactions with all other nodes in *R*. Since the only thing that the elements of *R* have in common is that they are TSS identified by the set of input SNPs, unexpectedly strong inter-relationships between elements of *R* are taken as indirect evidence of a relationship between the input SNPs themselves. The NDA score, *s*, is defined as the sum of –*log*_10_(*p*) values for interaction strength within the matrix.

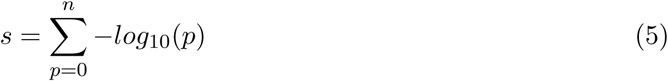

Raw *p*-values are calculated from the empirical distribution of values of *s* for 10000 permuted networks. The Benjamini-Hochberg method is used to estimate false discovery rate (*FDR*). Significant network density scores are taken as those with *FDR* < 0.05. In order to enable coexpression scores to be compared loosely between different analyses, each raw coexpression scores (*s*) is corrected by dividing by the total number of independent groups of regulatory regions included in each analysis, *n_r_es*, yeilding a corrected coexpression score, *ccs*:

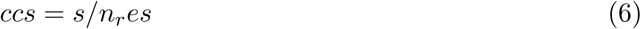

#### 2.6 Iterative recalculation

The node in the network with the highest NDA score has, by definition, numerous strong correlations with other nodes in the subset *R*. The NDA scores assigned to these other nodes are therefore inflated by their association with the stongest node. This inflation may reflect biological reality, since both TSS have a putative genetic association with the phenotype of interest, and both share strong links. However, there is a risk that TSS sharing a chance association with a strongly coexpressed TSS will be spuriously inflated to significance. For this reason, we have applied a stringent correction in order to ensure that we have confidence in each significantly coexpressed TSS independently of all TSS with stronger coexpression in the network: the NDA score for each TSS is calculated after removing all TSS with stronger NDA scores from the network.

#### 2.7 Input datasets

Of 267,225 robust promoters and enhancers identified by FANTOM5[6], 93,558 (50.6%) were promoters within 400 bases of the 5′ end of a known transcript model[6]. These were annotated with the name of the transcript. Alternative promoters were named in order of the highest transcriptional activity[6]. Where necessary, coordinates for GWAS SNPs (see 2.11 were translated to hg19 coordinates using LiftOver[9], or coordinates were obtained for SNP IDs from dbSNP[14] version 138.

#### 2.8 Permutations

A circular permutation method was devised to prevent systematic bias by maintaining the underlying structure of GWAS SNP data. The NDA score for a given regulatory region was compared with NDA scores obtained from randomly permuted subsets of genes to give an empirical *p*-value for coexpression. If permuted networks consist of randomly-selected regulatory regions, then this *p*-value quantifies coexpression alone (see 2.8.1); if the permuted networks are generated by mapping randomly-selected SNPs to regulatory regions, then the final *p*-value is a composite of two measures: coexpression, and the enrichment for true GWAS hits in regulatory sequence (see 2.8.2)).

##### 2.8.1 Pre-mapping permutations

Pre-mapping permutations use a random set of SNPs generated by rotation of the input set of SNPs, *K*, on a concatenated circular genome. The choice of background is critical - some more recent GWAS studies consider only a subset of variants with a high probability of association with a given trait, such as the immunochip[16] or the metabochip[17]. In the present analyses, background data were chosen to reflect as accurately as possible the pool of variants included in the original study. For this reason, results are presented only for phenotypes for which the the entire summary dataset was available, including a *p*-value for every SNP, so that the background used to generate permuted networks is exactly the same background from which the real dataset is drawn.

##### 2.8.2 Post-mapping permutations

In order to quantify the effect of coexpression alone (i.e. eliminating the inflation of NDA scores that occurs due to enrichment of trait-associated SNPs in regulatory regions), permuted networks were generated after mapping to TSS regions. Let *A* be the whole set of FANTOM5 TSS. Post-mapping permutations select a random subset of *A* in a similar circular manner, by randomly displacing the members of the set *R* (Equation 3) by a random number of places on the list. Where the displacement pushes members of *R* off the end of the list, they are re-entered at the beginning.

This process generates a pool of variants that are likely to be grouped in a similar distribution on the genome as the input set. If the input set contains a large group of TSS regions in close proximity on the genome, it is likely that this group of TSS regions will be joined as a single unit (see above) for analysis. During generation of permutations, the same number of consecutive TSS regions elsewhere on the genome may not be in sufficient proximity (and expression correlation) to be grouped together. This would create extra network nodes, falsely inflating the NDA scores in the permuted sets. In order to mitigate against this, those TSS from each permutation that do not conform to the input set distribution are re-entered into a further circular permutation until an identical distribution is found. If no matching grouping is found after 8 repeat permutations, additional regulatory regions are added from consecutive positions above and below whichever group is nearest in size to the relevant group in the original input dataset.

**Supplementary Figure 1:**
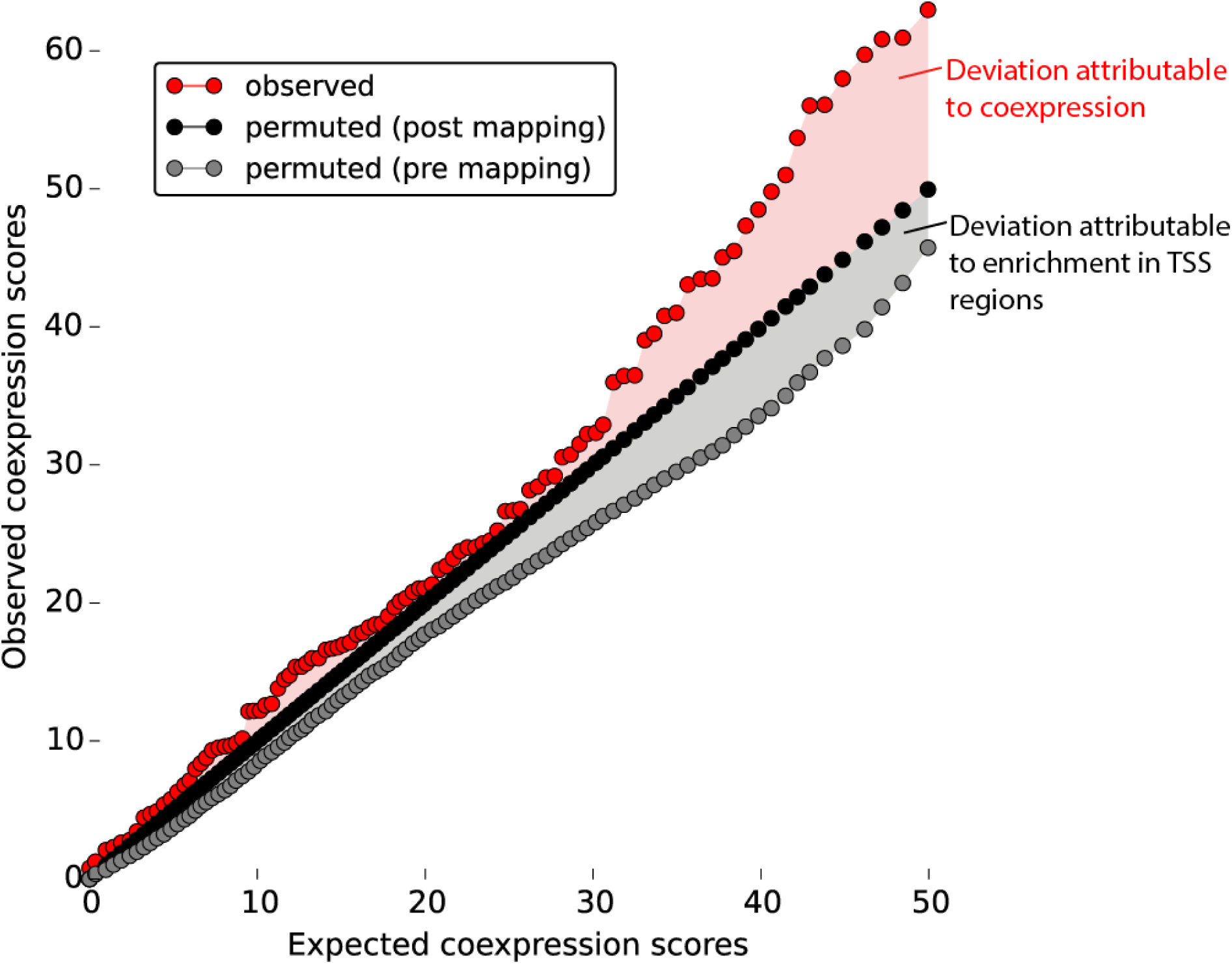
Distribution of observed NDA scores for Crohn’s disease, and expected NDA scores from pre and post-mapping permutations.

The difference between the distributions of NDA scores derived from pre and post-mapping permutations reveals the different components of the measure. When compared to a random pool of SNPs (pre-mapping permutations), two factors inflate the NDA scores for real GWAS data: firstly, more regulatory regions are identified because true GWAS hits are enriched within regulatory regions; secondly, the coexpression signal itself is greater for real data. In contrast, post-mapping permutations have precisely the same number of regulatory regions included as the real dataset, so there is no component of inflation due to enrichment in regulatory regions. The effects of these different components are shown in Figure 1, which reveals the NDA score to be a composite measure of both signals.

False discovery rates (FDR) are calculated using the Benjamini-Hochberg method[2].

#### 2.9 Choice of samples and regulatory regions

The enrichment for GWAS hits from a pooled resource comprising the NCBI GWAS catalog and the GWASdb database (observed *SNPs.Mb*^−1^: expected *SNPs.Mb*^−1^) was quantified at increasing search window sizes upstream and downstream from the transcription start site (TSS). A table of GWAS hits for a broad range of phenotypes was obtained from the NCBI GWAS catalog[8] and from a larger, less selective catalog of GWAS *p*-values meeting permissive criteria for genome-wide significance, GWASdb[13]. The GWASdb dataset is less fastidiously curated than the NCBI GWAS catalog, but contains a much greater range of SNPs since it does not restrict inclusion to the strongest associations, or to putative causative variants. Since both databases are limited by the variation in reporting, and quality, of the original GWAS studies from which data are drawn, this analysis was restricted to variants meeting genome-wide significance at a widely-accepted threshold (*p* < 5 × 10^−8^). These catalogues were combined and filtered to remove duplicate entries. Data were obtained from:

- NHGRI GWAS catalog, June 2014 http://www.genome.gov/gwastudies
- GWASdb2, June 2014 update ftp://jjwanglab.org/GWASdb/20140629/gwasdb_20140629_snp_trait.gz

Overlapping phenotypes, such as “urate” and “uric acid” were manually merged as shown in SF2_phenotype_matching.txt. Phenotypes that were considered to be too broad to be informative were excluded, as were those that were not related to human disease. A complete table of phenotypes in GWASdb and NCBI GWAS catalog, showing mergers and inclusion/exclusion in the present work, is provided in a supplementary file (SF2_phenotype_matching.txt).

The coexpression signal obtained for the test input set was evaluated using different subsets of FANTOM5 samples (cell lines, timecourses following a perturbation in primary cells or selected cell lines, tissue samples, primary cells, or various combinations of these), and different types of regulatory region (enhancers, promoters assigned to annotated genes, other promoters, or all regulatory regions combined)(Supplementary Figure 3). A weak signal for coexpression is seen in cell lines, but the addition of cell lines to the combined sample set of primary cells, timecourses and tissues did not improve the coexpression signal seen for any subset of regulatory regions. The strongest coexpression is seen in the combined sample set. A “minimal detail” sample set was also tested, comprising a single average value for each of the timecourses, primary cell types and tissue types, and excluding data from unstimulated cell lines. The complete dataset, including all cell types and tissues, provided the strongest signal, demonstrating that there is additional biologically-relevant information contained in the expression profiles from all sample subsets (Supplementary Figure 3).

#### 2.10 Anti-correlation

Strong anti-correlation between pairs of TSS associated with the same phenotype may have biological importance, such as down-regulation at one TSS but expression at another, or negative regulation of a signalling pathway on which expression of a TSS is dependent. For this reason, anti-correlations may improve detection of true associations in this analysis. However, in order to confer an overall improvement on the performance of the algorithm, true inverse expression relationships between phenotype-associated TSS would need to be sufficiently common to overcome the noise added by incorporating all strong anti-correlations into the NDA score.Anti-correlations do not contribute any net improvement to the NDA scores for a training set (Crohn’s disease, 50% of all SNPs, chosen at random), and were therefore excluded.

#### 2.11 GWAS data sources

Full GWAS or meta-analysis data, reporting every SNP genotyped or imputed in a given study, are required in order to permute subsets against the appropriate background for a given study (see 2.8). These were obtained from the following sources:

- Crohn’s disease[7] summary *p*-values were obtained from the International Inflammatory Bowel Disease Genetics Consortium ftp://ftp.sanger.ac.uk/pub4/ibdgenetics/cd–meta.txt.gz
- Ulcerative colitis[1] summary *p*-values were obtained from the International Inflammatory Bowel Disease Genetics Consortium ftp://ftp.sanger.ac.uk/pub4/ibdgenetics/ucmeta–sumstats.txt.gz
- Summary *p*-values for human height[3] were obtained from the GIANT consortium https://www.broadinstitute.org/collaboration/giant/images/4/47/GIANT_HEIGHT_LangoAllen2010_publicrelease_HapMapCeuFreq.txt
- Summary *p*-values for total cholesterol, LDL cholesterol, HDL cholesterol and triglycerides[5] were obtained from the Global Lipids Consortium http://csg.sph.umich.edu/abecasis/public/lipids2013/
- Summary *p*-values for systolic and diastolic blood pressure. [15] were obtained from the International Consortium on Blood Pressure study http://www.georgehretlab.org/icbp_088023401234-9812599.html

A permissive threshold for trait association of *p* < 5 × 10^−6^ was used for whole GWAS / metaanalysis coexpression analyses.

### 3 Cell type specificity

In order to better understand the pathophysiological implications of disease variants in regulatory regions, we sought to identify whether these regions exhibit unexpectedly specific expression in any given cell types or tissue samples. In order to reduce noise, technical and biological replicates were averaged for this and subsequent analyses. The full table of samples in FANTOM5, showing which samples were averaged as technical replicates, and which were excluded, is in supplementary table (SF2_phenotype_matching.txt).

For a given trait, we took the subset of regulatory regions for which a significant coexpression pattern was detected for that trait (coexpression *FDR* ≤ 0.05). For each regulatory region, we created a list of all cell types in which that region was active, ranked by expression level. We then combined the cell type lists for each regulatory region using a robust rank aggregation (RRA)[11].

There are several possible sources of bias in this raw measurement. For example, some cell types have more cell-type specific transcriptional activity, perhaps because these cell types fulfil a specialised role; other cell types are particularly well-represented in the FANTOM5 samples. We therefore controlled for the probability that a given cell type would be highly ranked in the initial RRA analysis, by permuting RRA results for at least 100,000 random selections of n regulatory regions. We then calculated the empirical *p*-value for a each cell type, i.e. the probability that this cell type would be assigined a raw RRA *p*-value at least as strong by random chance. We then corrected for multiple comparisons using the Benjamini-Hochberg method to estimate false discovery rate (*FDR*).

### 4 Code availability

Computer code required to run the NDA method, specifically for the detection of coexpression in FANTOM5 regulatory regions, can be obtained from https://github.com/baillielab/coexpression/

**Supplementary Figure 2:**
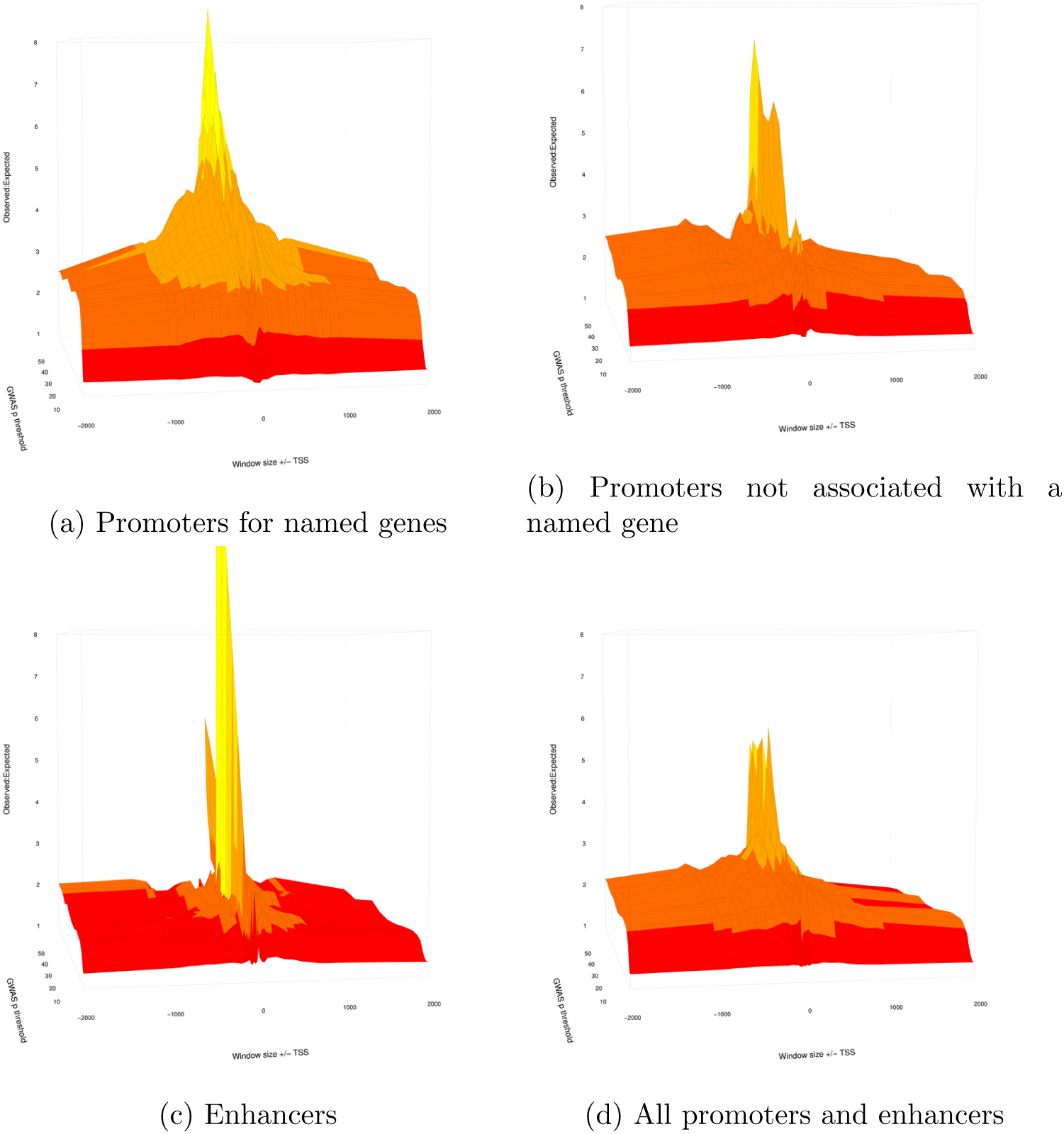
Enrichment for GWAS hits at increasing distances above and below TSS.

**Supplementary Figure 3:**
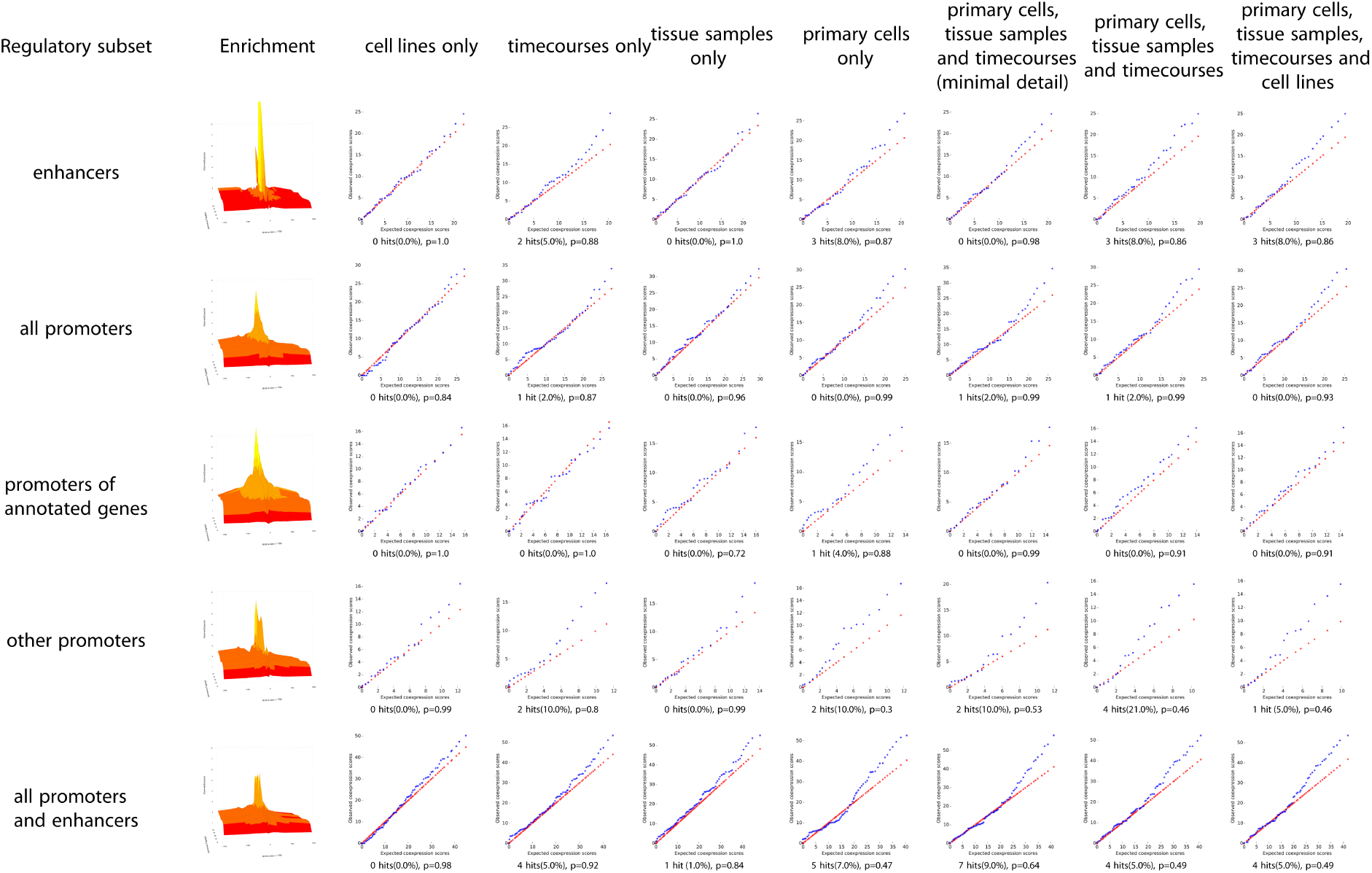
Change in coexpression signal using different subsets of the FANTOM5 dataset, using the Crohn’s disease GWAS as the input set. Enrichment column shows a miniature graph depicting the enrichment (observed *SNPs.Mb*^−1^: expected *SNPs.Mb*^−1^) at increasing search window sizes upstream and downstream from the transcription start site (TSS). Other columns show Q:Q plots of observed:expected NDA scores obtained using a given subset of samples (see SF3_sample_averaging.xlsx for full description of each subset). Rows indicate the subset of regulatory regions used in each analysis. Percentage of significantly coexpressed entities (hits, *FDR* < 0.05) and *p*-value (Kolmogorov-Smirnov test) comparing observed (blue) and expected (red) distributions are shown below each plot.

